# Metabolism of glucose and glutamine is critical for skeletal muscle stem cell activation

**DOI:** 10.1101/2020.07.28.225847

**Authors:** Sanjana Ahsan, Manmeet H. Raval, Maxwell Ederer, Rajiv Tiwari, Andrew Chareunsouk, Joseph T. Rodgers

**Affiliations:** The Edythe and Eli Broad Center for Regenerative Medicine and Stem Cell Research, Department of Stem Cell Biology and Regenerative Medicine, Keck School of Medicine, University of Southern California, 90033

## Abstract

Injury to muscle tissue induces the resident, quiescent, skeletal muscle stem cells (MuSCs) to activate - to exit quiescence and enter the cell cycle. Previous work has shown that MuSC activation is associated with significant metabolic changes, however the substrates that MuSCs consume to support activation are poorly understood. Here, we show that MuSCs generate the majority of their energy through mitochondrial respiration, and that oxidative phosphorylation is required for MuSC activation. Furthermore, we have found that while glucose, glutamine, and fatty acids all significantly, and roughly equally, contribute to ATP production in MuSCs during activation, they do not have equal functional role in the dynamics of MuSC activation. Pharmacologic suppression of glycolysis, using 2-deoxy-D-glucose, or glutaminolysis, using BPTES, significantly impairs MuSC cell cycle entry. However, etomoxir-mediated inhibition of mitochondrial fatty acid transport has minimal effect on MuSC cell cycle progression. Our findings suggest that apart from their roles in fueling ATP production by the mitochondria, glucose and glutamine may generate metabolic intermediates needed for MuSC activation.

## INTRODUCTION

Skeletal muscle has a robust capacity to regenerate after injury. This regenerative ability largely depends on the function of its tissue resident muscle stem cells (MuSCs), also known as satellite cells. Under homeostatic conditions, MuSCs exist in a quiescent state characterized by small size, low transcription, and low metabolic activity^1–5^ When muscle is injured, signals from the damaged tissue environment induce MuSCs to activate, i.e. to exit the quiescent state, enter the cell cycle, and proliferate. MuSC activation has been found to correlate with increases in cellular and metabolic activity^1,5^. In addition, previous work has shown that activation is a critical step in muscle repair, in that defects in activation contribute to impairments in regeneration. Conversely, MuSCs with improved activation function contribute to improvements in muscle regeneration^1,6,7^.

Metabolic signals are known to be integrally linked with MuSC activation and muscle regeneration. Previous work has shown that MuSCs from mice that were subjected to restricted diet had increases in mitochondrial function which correlated with improvements in muscle regeneration^8^. Additionally, deficiency in SIRT1 activity leads to defects in autophagic flux needed to meet the bioenergetic demands of MuSC activation, and consequently delays MuSC activation and muscle regeneration ^3,5^. Furthermore, loss of function of pyruvate dehydrogenase kinase (PDK) 2 and PDK4 in MuSCs contributes to dysregulation of glucose metabolism during activation and is associated with impaired muscle regeneration^2^. While this work ha shown the important role of metabolism in MuSC activation, there have been no systematic investigations of metabolic substrate utilization by MuSCs. Here, we report that during activation, MuSCs display a profound increase in ATP production that is essentially entirely produced by the mitochondria. We find that glucose, glutamine, and fatty acids contribute roughly equally to ATP production during activation. However, we find that MuSC cell cycle entry is specifically dependent upon glucose and glutamine oxidation. Collectively, these data provide a new detailed analysis of substrate metabolism during MuSC activation.

## RESULTS

### Cellular activity significantly increases in MuSCs during activation

To gain a more detailed understanding of the metabolic function of MuSCs during activation, we employed an *ex vivo* model of isolation-induced activation^1,3^. Immediately after dissociation from the hind limb muscle of adult mice (3.5 - 7 months old) and FACS-mediated purification, we found that essentially no freshly isolated (FI) MuSCs were positive for phosphorylated Ser 807/811-RB (pRB), a marker of cells that have exited the G1 phase of the cell cycle, or incorporated EdU nucleotide, suggesting that the vast majority of these cells were in early G1 (Figures 1A and 1B). After culturing for 24 and 40 hours post-isolation (hpi), the number of MuSCs positive for these markers progressively increased, demonstrating that MuSCs enter the cell cycle following isolation. We used time-lapse microscopy to continuously monitor cultures of MuSCs for the first 90 hours after isolation and found that MuSCs require a median of 48 hours to complete cytokinesis after isolation (Figure 1C), consistent with previous work ^1,9^.

**Figure 1.**
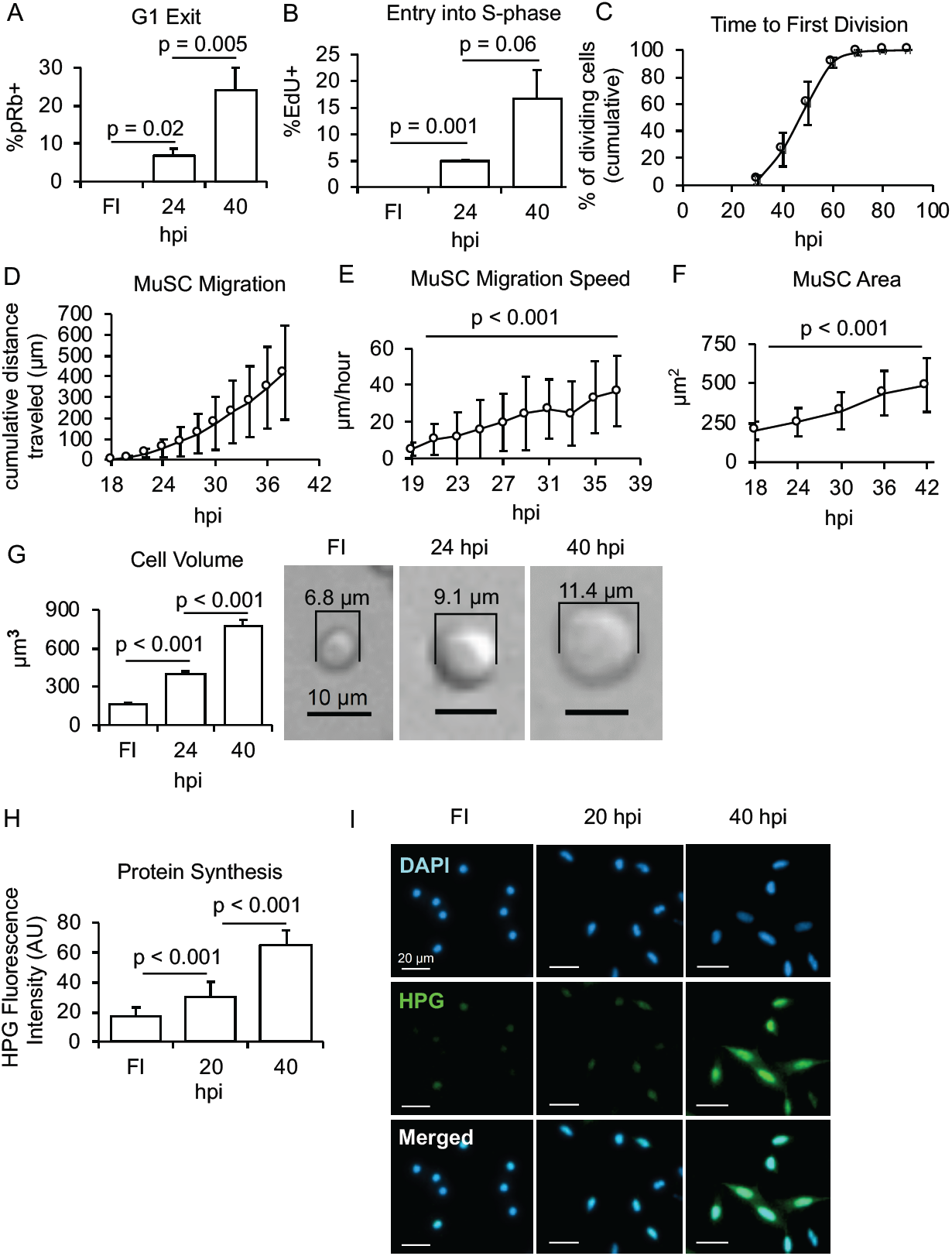
Cellular activity increases in MuSCs during activation. **(A & B)** Proportion of MuSCs positive for pRb (A) and EdU (B) when they are freshly isolated (FI), 24, and 40 hours post-isolation (n = 3 - 4). **(C)** Cumulative proportion of dividing MuSCs that have completed the first division following isolation (n = 5). **(D)** Cumulative distance traveled by MuSCs from 18 to 38 hpi (n = 30 cells). **(E)** MuSC migration speed between 19 to 37 hpi (n = 30 cells). Significance was calculated between 19 and 37 hpi. **(F)** Average MuSC area between 18 to 42 hpi (n = 40 cells). Significance was calculated between 18 and 42 hpi. **(G)** Average cell volume of MuSCs at 0 (n = 4), 24 and 40 (n = 3) hours after isolation (left) and representative images (right). **(H)** Protein translation in MuSCs detected through incorporation of HPG into newly synthesized proteins at 0 (n = 87 cells), 20 (n = 101 cells), and 40 (n = 103 cells) hpi. **(I)** Immunofluorescence images of MuSCs stained for DAPI and HPG.

Readily apparent in our time-lapse microscopy experiments was that MuSCs displayed tremendous changes in cell migration and size during activation (Supplemental Video 1). We found that MuSCs migrated a cumulative distance of 419 µm in the first 38 hours after plating (Figure 1D). Interestingly, we noticed that the distance versus time graph (Figure 1D) of MuSC migration was convex, suggesting that the speed of MuSC migration was also increasing. We analyzed the average speed of MuSC migration by measuring distance MuSCs traveled between each 2-hour time window, and found that the speed at which MuSCs move also increases during activation, from 5.1 µm/hour at 19 hpi to 36.9 µm/hour at 37 hpi (Figure 1E). Similarly, we found that the 2-dimensional area of MuSCs increases during activation from 199.9 µm^**2**^ at 18 hpi to 489 µm^**2**^ at 50 hpi (Figure 1F).

To determine if the changes in cell area reflected changes in cell size, we approximated cell volume from diameter measurements of trypsinized MuSCs, which are roughly spherical, at various time points during activation. We found that in the first 40 hpi, cell volume increases by nearly five-fold, from 164 µm^**3**^ to 774 µm^**3**^ (Figure 1G). To determine if these changes in cell size reflected cell growth or anabolism, we measured protein synthesis rates. To do this, we pulsed MuSCs with L-homopropargylglycine (HPG), an amino acid analog that incorporates into nascent protein synthesis and can be detected using Click-It chemistry (Invitrogen), to measure protein translation rates. We found that after a 2-hour pulse with HPG, 40 hpi MuSCs incorporate about four-fold more HPG than freshly isolated MuSCs, suggesting that protein translation rates increase during activation (Figures 1H and 1I). Collectively, these data show that MuSCs display tremendous changes in cell size and activity as they enter and progress through the cell cycle following isolation.

### MuSC activation requires ATP production from the mitochondria

These dramatic increases in cellular activity during activation suggested that cellular metabolism would also need to significantly change to fuel these processes ^10^. To investigate this, we performed a series of live cell metabolic flux analyses on MuSCs using a Seahorse bioanalyzer. We found that MuSCs display a dramatic overall increase in oxygen consumption rate (OCR) in the first 48 hours after isolation (Figure 2A). After subtracting the contribution of non-mitochondrial OCR, we found that mitochondrial dependent OCR increases by roughly 16-fold over 48 hours after isolation (Figure 2B). Additionally, we measured the extracellular acidification rate (ECAR), or the rate at which metabolic activity in MuSCs acidifies the media. Similar to OCR, we found that MuSC ECAR increases by ten-fold during the first 48 hours after isolation (Figure 2C). These results show that there is a dramatic increase in MuSC metabolic activity during activation.

**Figure 2.**
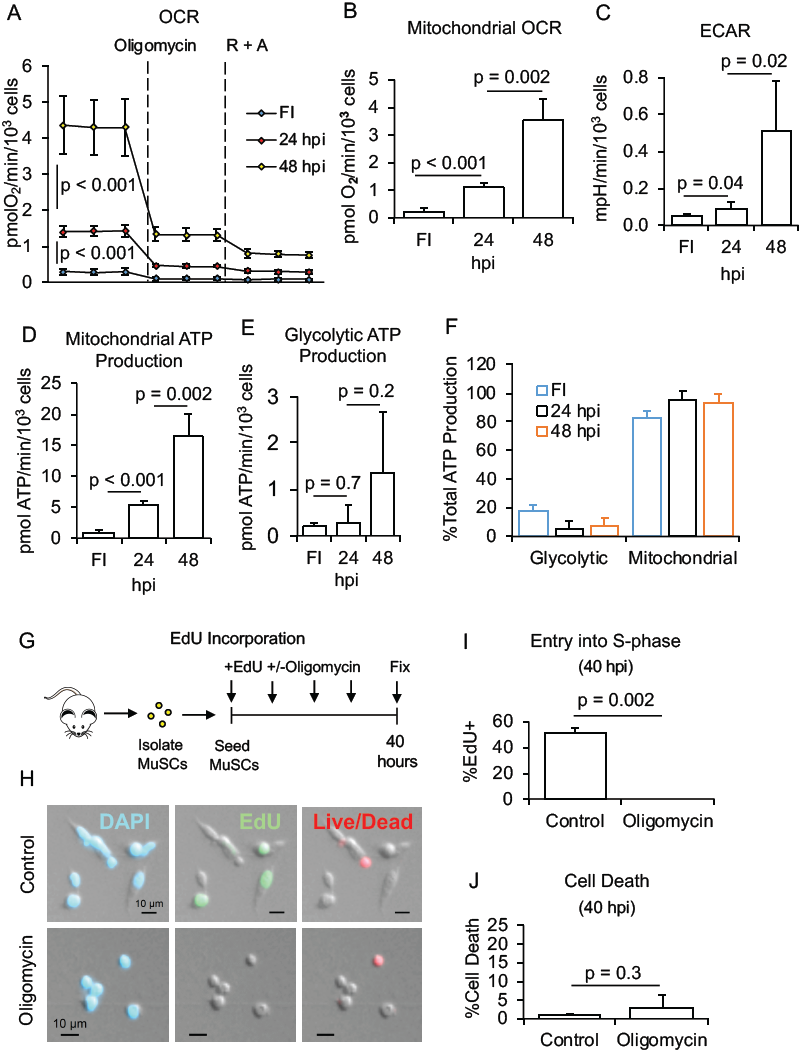
MuSCs require mitochondrial metabolism to activate. **(A)** Changes in basal OCR in MuSCs after inhibition of ATP synthase using oligomycin, followed by inhibition of Complex I and III of the electron transport chain with rotenone and antimycin A (R + A) (n = 5). **(B)** Total OCR derived from mitochondria at 0 (n = 4), 24 and 48 hpi (n = 5). **(C)** ECAR at 0, 24, and 48 hpi (n = 5). **(D)** Rate of mitochondrial ATP production in MuSCs at 0, 24, and 48 hpi (n = 5). **(E)** Rate of glycolytic ATP production in MuSCs at 0, 24, and 48 hpi (n = 5). **(F)** Relative proportion of MuSC total cellular ATP production derived from mitochondrial respiration and glycolysis at 0, 24, and 48 hpi (n = 5). **(G)** Experimental schematic of the EdU incorporation assay in MuSCs treated with or without oligomycin. **(H)** Immunofluorescence images of MuSCs stained for DAPI, EdU, and Live/Dead cell viability stain. **(I)** Proportion of control and oligomycin-treated MuSCs that have incorporated EdU at 40 hpi (n = 3). **(J)** Cell death in MuSCs cultured with or without oligomycin for 40 hpi (n = 3).

Previous work has shown that OCR and ECAR measurements can be used to calculate ATP production rates from glycolysis and the mitochondria ^11,12^. Our analysis showed that ATP production by the mitochondria displays very dramatic and significant increase in the 48 hours after isolation (Figure 2D). ATP production rate from glycolysis also displayed a similar trend of increase, but was not statistically significant (Figure 2E). Most interestingly, when we directly compared ATP production rates from the mitochondria and glycolysis, our measurements showed that the majority, ∼80%, of cellular ATP production comes from the mitochondria (Figure 2F). The relative contributions of glycolytic and mitochondrial ATP production did not change over the course of our measures, despite the strong increase in magnitude. These results show that majority of ATP produced in MuSCs during activation is from mitochondria.

The dramatic increase in mitochondrial ATP production rate suggested to us that mitochondrial metabolism plays a significant role in MuSC activation. To determine the requirement of mitochondrial ATP production during activation, we cultured cells in oligomycin, an inhibitor of the mitochondrial ATP synthase, for 40 hours post-isolation (Figure 2G). Treatment of cells with oligomycin dramatically decreases OCR due to feedback caused by the inability of cells to dissipate the mitochondrial proton gradient via ATP synthase (Figure 2A) ^11^. Interestingly, we found no obvious changes in cell death in MuSCs cultured in oligomycin for 40 hours (Figures 2H and 2J). However, we observed very strong changes in EdU incorporation: essentially zero MuSCs treated with oligomycin incorporated EdU in the first 40 hpi (Figures 2H and 2I). Our findings show that inhibition of mitochondrial ATP synthesis nearly completely blocks the ability of MuSCs to enter the S-phase of the cell cycle. These results suggest that mitochondrial ATP production is necessary for activation.

### Glucose oxidation is critical for MuSC activation

Given that mitochondrial ATP production is necessary for MuSC activation, we next investigated which metabolic substrates were consumed in activating MuSCs. Three major substrates that cells oxidize to generate ATP and biosynthetic intermediates are glucose, glutamine, and fatty acids ^2^. To investigate the contribution of glucose to MuSC activation, we treated cells with 2-deoxy-D-glucose (2-DG), an inhibitor of glycolysis, and examined the change in metabolic flux ^13^. We found that MuSCs treated with 2-DG displayed significantly reduced levels of total OCR, OCR-linked to ATP production (OCR_ATP_), and ECAR (Figures 3A-C). Using the measurements from control and 2-DG treated MuSCs, we calculated mitochondrial ATP synthesized from oxidation of glucose versus non-glucose substrates and found that both increased roughly three-fold from 24 to 48 hpi (Figure 3D). When we normalized to total mitochondrial ATP production (glucose + non-glucose), we found that glucose contributes to 29.6% of ATP generated by mitochondria at 24 hpi and 30.2% at 48 hpi (Figure 3E).

**Figure 3.**
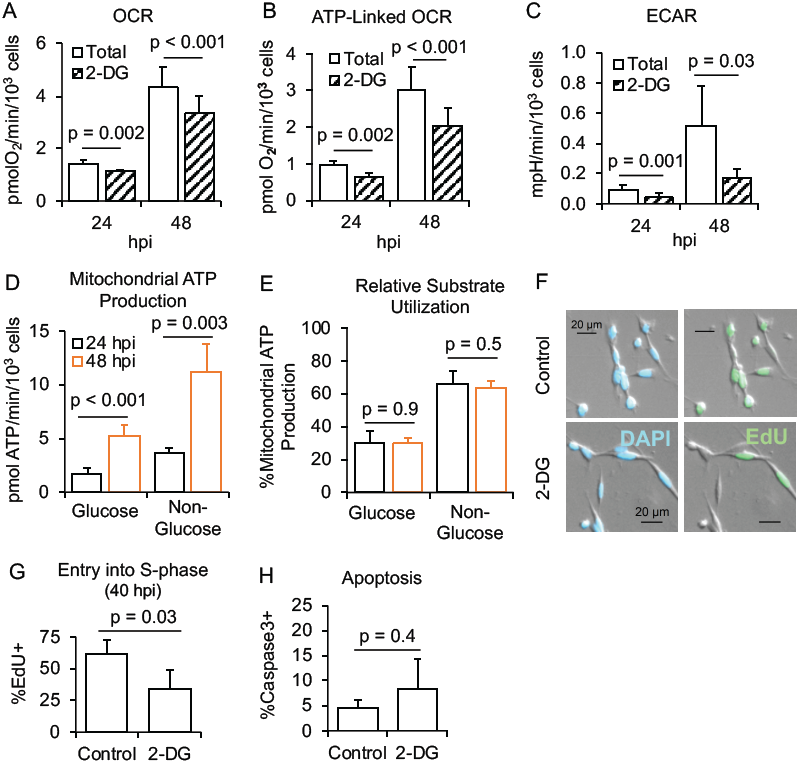
Glucose metabolism plays an important role in MuSC activation. **(A & B)** Changes in total (A) and ATP-linked (B) oxygen consumption rate induced by injection of 2-DG at 24 and 48 hpi (n = 5). **(C)** Total ECAR before and after 2-DG injection at 24 and 48 hpi (n = 5). **(D)** Mitochondrial ATP production derived from glucose and non-glucose substrates at 24 and 48 hpi (n = 5). **(E)** Proportion of MuSC mitochondrial ATP production derived from glucose and non-glucose substrates at 24 and 48 hpi (n = 5). **(F)** Immunofluorescence images of control and 2-DG treated MuSCs stained for DAPI and EdU at 40 hpi. **(G & H)** Proportion of control and 2-DG treated MuSCs that are positive for EdU (G; n = 4) and Caspase 3 (H; n = 4 for control, n = 3 for 2-DG) at 40 hpi.

Our finding that glucose accounts for nearly one-third of mitochondrial ATP production suggested that glucose plays an important role in MuSC activation. We therefore tested the ability of MuSCs to activate under suppression of glycolytic metabolism. To do this, we cultured the cells in 2-DG for 40 hours following isolation and measured EdU incorporation. We found that compared to MuSCs cultured under control conditions, a smaller proportion of MuSCs activate under 2-DG treatment (Figures 3F and 3G). Detection of caspase 3-positive MuSCs showed that inhibition of glycolysis does not significantly change the proportion of cells undergoing apoptosis (Figure 3H). Thus, in contrast to the near complete block of activation by oligomycin, the data on 2-DG indicate that glycolysis has an important role, but is not absolutely required for MuSC activation.

### Glutamine is a major metabolic substrate during MuSC activation

Glutamine is another major metabolic substrate that is oxidized by the mitochondria via the TCA cycle. To examine the contribution of glutamine metabolism in MuSC activation, we treated cells with Bis-2-(5-phenylacetamido-1, 3, 4-thiadiazol-2-yl)ethyl sulfide (BPTES), a small molecule which inhibits glutaminase, which converts glutamine to glutamate ^14^. Glutamate is a precursor for the TCA cycle intermediate α-ketoglutarate; therefore, we expected BPTES treatment to suppress mitochondrial oxygen consumption in activating MuSCs ^15^. Interestingly, we found that culturing cells with BPTES for 24 hours did not have a significant effect on total OCR or OCR_**ATP**_ (Figures 4A - C). However, BPTES had a very pronounced effect at reducing OCR, and consequently mitochondrial ATP production rate, at 48 hpi (Figure 4A - C, 4E). BPTES did not have a significant effect on ECAR at either time point (Figure 4D). By comparing the measurements from control and BPTES treated cells, we calculated the proportion of mitochondrial ATP from glutamine and non-glutamine substrates and found that glutamine accounts for 8% of mitochondrial ATP production at 24 hpi, and 38% at 48 hpi (Figure 4F).

**Figure 4.**
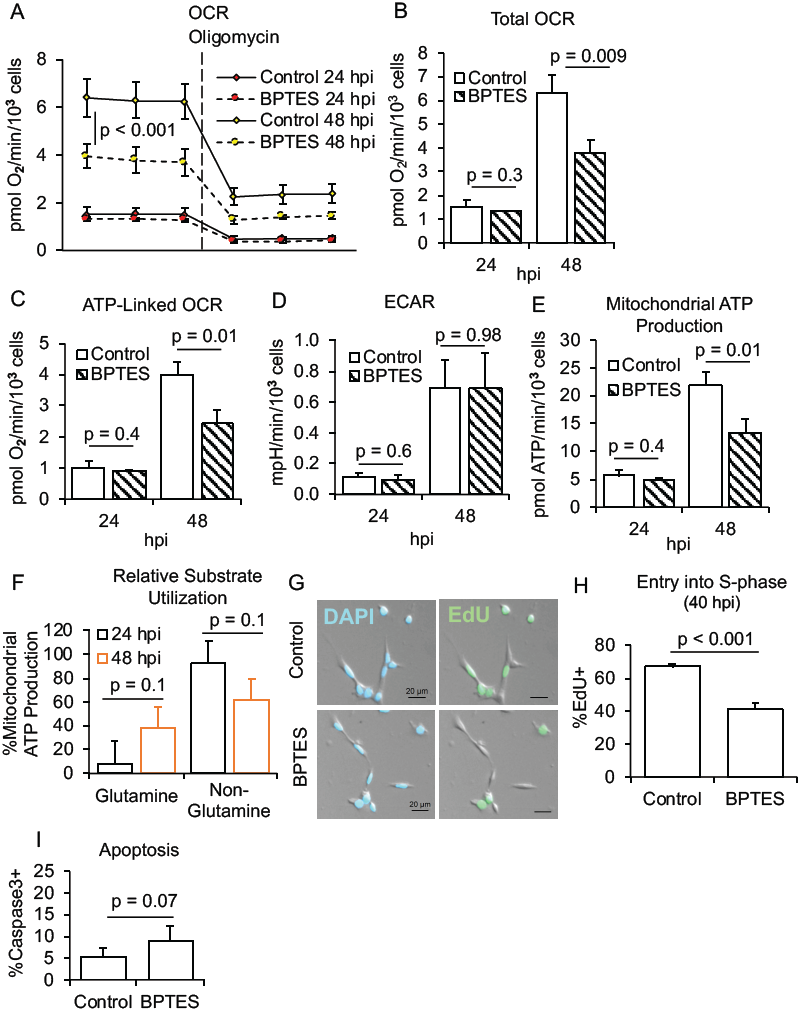
Glutamine metabolism is important but not necessary for MuSC activation. **(A)** Changes in OCR induced by oligomycin injection in MuSCs cultured with or without BPTES for 24 and 48 hpi (n = 3). **(B - D)** Total (B) and ATP-linked (C) OCR and ECAR (D) in MuSCs cultured with or without BPTES for 24 and 48 hpi (n = 3). **(E)** Mitochondrial ATP production in control and BPTES-treated MuSCs (n = 3). **(F)** Proportion of MuSC mitochondrial ATP production derived from glutamine and non-glutamine substrates at 24 and 48 hpi (n = 3). **(G)** Immunofluorescence images of control and BPTES-treated MuSCs stained for DAPI and EdU at 40 hpi. **(H & I)** Proportion of control and BPTES-treated MuSCs that are positive for EdU (G; n = 4) and Caspase 3 (H; n = 5 for control, n = 4 for BPTES) at 40 hpi. Data are presented as mean ± standard deviation.

Although exposure to BPTES for 24 hours did not have a significant effect on MuSC mitochondrial ATP production, the magnitude of effect observed under 48-hour BPTES treatment was similar to that observed under inhibition of glycolysis. Based on these results, we predicted that inhibition of glutamine metabolism would impair MuSC activation similar to the effect of 2-DG. To determine the role of glutaminolysis in activation, we cultured MuSCs for 40 hours with or without BPTES treatment and measured EdU incorporation. As expected, fewer MuSCs had incorporated EdU at 40 hpi in cultures treated with BPTES than control (Figures 4G and 4H). BPTES treatment induced a modest increase in apoptosis in MuSCs (Figure 4I). These results suggest that suppression of glutaminolysis slows down MuSC activation, and that similar to glycolysis, glutaminolysis is important but not necessary for activation.

### Fatty acid metabolism is dispensable for MuSC activation

Finally, we investigated the role of fatty acid β-oxidation in MuSC activation. To do this, we treated cells with etomoxir, an inhibitor of carnitine palmitoyltransferase I, to block the transport of fatty acids into the mitochondria ^16^. Similar to BPTES, treatment with etomoxir for 24 hours did not have a statistically significant effect on OCR or ECAR (Figures 5A-D). However, after 48 hours of etomoxir treatment, total OCR, OCR_ATP_, and mitochondrial ATP production rates were significantly reduced compared to control (Figures 5A-C, 5E). We determined the relative contribution of fatty acids to mitochondrial ATP production and found that there was a trend of increase from 26.2% at 24 hpi to 38.8% at 48 hpi (Figure 5F). These results suggest that fatty acids may be an important metabolic substrate during MuSC activation.

**Figure 5.**
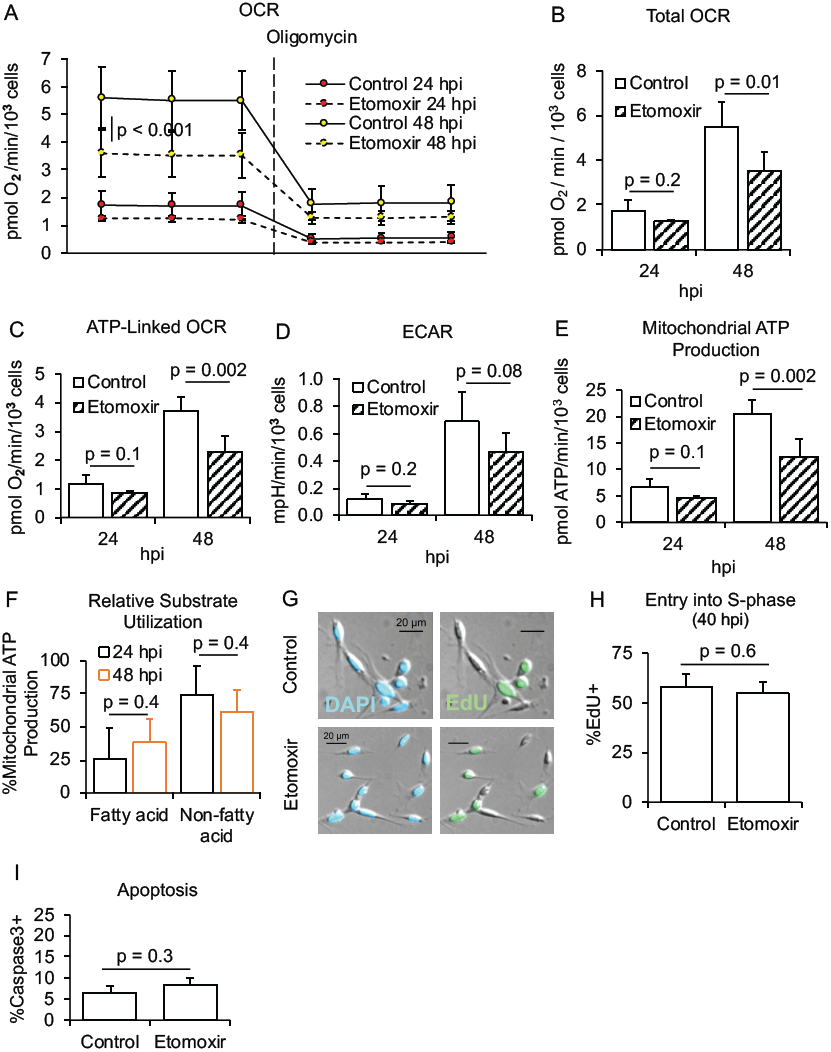
Fatty acid metabolism is not necessary for MuSC activation. Fatty acid metabolism is not required for MuSC activation. **(A)** Changes in OCR induced by oligomycin injection in MuSCs treated with or without etomoxir for 24 (n = 3) and 48 (n = 5) hpi. **(B - D)** Total (B) and ATP-linked (C) OCR and ECAR (D) in MuSCs cultured with or without etomoxir for 24 (n = 3) and 48 (n = 5) hpi. **(E)** Mitochondrial ATP production in control and etomoxir-treated MuSCs at 24 (n = 3) and 48 (n = 5) hpi. **(F)** Proportion of MuSC mitochondrial ATP production derived from fatty acid and non-fatty acid substrates at 24 (n = 3) and 48 (n = 5) hpi. **(G)** Immunofluorescence images of control and etomoxir-treated MuSCs stained for DAPI and EdU at 40 hpi. **(H & I)** Proportion of control and etomoxir-treated MuSCs that are positive for EdU (H) and Caspase 3 (I) at 40 hpi (n = 3).

Based on the reduced ability of MuSCs to enter the S-phase under inhibition of glucose and glutamine metabolism, we predicted that etomoxir, by blocking fatty acid entry into the mitochondria, would also slow down MuSC activation. Surprisingly, however, MuSCs treated with etomoxir for 40 hours had the same level of EdU incorporation as control (Figures 5G and 5H). We did not observe changes in apoptosis under etomoxir exposure (Figure 5I). Collectively, these data suggest that while MuSCs utilize fatty acids as a fuel source during activation, fatty acids are not critical for the entry of MuSCs into S-phase.

## DISCUSSION

Metabolism has been shown to play a crucial role in the function of many tissue resident stem cells ^17^. Specifically in the context of MuSCs, recent work has shown that MuSCs display an increase in glycolytic metabolism during activation and this is important for muscle regeneration ^2,3^. We similarly observed that rates of glycolysis increase during activation (Figures 2C and 2E). Interestingly, we found that the ATP production from glycolysis actually decreases, relative to ATP production from the mitochondria (Figure 2F). Our data show that glycolysis is an important pathway during MuSC activation, as inhibition of glycolysis by 2-DG significantly blunts entry of MuSCs into the S-phase of the cell cycle (Figure 3G). However, our data suggest that a function of glycolysis during activation is to provide substrates for further oxidation in the mitochondria. Our findings are consistent with recent work showing that MuSC-specific deletion of the transcription factor Ying Yang 1 (YY1), which promotes mitochondrial metabolism ^18–20^, contributes to reduction of mitochondrial respiration and impairs the ability of MuSCs to enter the cell cycle ^21^.

Our analysis of MuSC substrate metabolism showed that glucose, glutamine, and fatty acids can contribute to mitochondrial ATP synthesis at similar levels in activating MuSCs (Figure 6). All three metabolic substrates, however, were not equally important for MuSCs to enter the S-phase of the cell cycle. While suppression of glucose and glutamine metabolism delayed entry into S-phase, inhibiting fatty acid metabolism had no effect. One possible explanation for the lack of effect of etomoxir on S-phase entry is that MuSCs are able to compensate by increasing other metabolic pathways. We analyzed ECAR and found that it did not significantly change in response to etomoxir (Figure 5D), suggesting that glycolysis rates did not increase to compensate for inhibition of fatty acid oxidation. As mitochondrial ATP production rates only displayed a modest, and not statistically significant decrease in response to etomoxir at 24 hpi, this may suggest that MuSCs compensate by increasing glutamine oxidation. Further work is needed to test the effect of etomoxir treatment on MuSC glutamine metabolism. An alternative explanation is that fatty acid oxidation may not have an important role in fueling the early phases of MuSC cell cycle entry. Indeed, the strongest effect of etomoxir is after 48 hours of activation, a stage when the most MuSCs have already entered and completed the first cell cycle after isolation ^1^. This suggests that fatty acid oxidation may have an important role in the daughter cells from MuSC division. These data show that while fatty acid metabolism contributes to energy production in activating MuSCs, it is not necessary for the cells to activate. Inhibition of individual metabolic pathways for each substrate only partially reduced mitochondrial metabolism; it is likely that combined suppression of all three pathways will block mitochondrial metabolism and cell cycle entry, similar to oligomycin.

**Figure 6.**
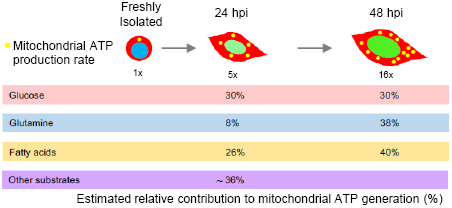
Summary of MuSC metabolic substrate utilization. Summary of MuSC metabolic substrate utilization during activation. Mitochondrial ATP production rates increase 16-fold from freshly isolated to 48 hours post isolation. Oxidation of glucose by the mitochondrial accounts for ∼30% of the ATP generated at 24 and 48 hpi. The contribution of glutamine and fatty acid oxidation to mitochondrial ATP production increases from 24 to 48 hours. At 48 hpi the majority of ATP production by the mitochondrial is from glucose, glutamine, and fatty acid oxidation. However, at 24 hpi roughly 36% of mitochondrial ATP production is from substrates other than glucose, glutamine, and fatty acids.

Although glucose, glutamine, and fatty acids are able to support ATP synthesis in MuSCs, only inhibition of glucose and glutamine metabolism delays MuSC activation. This indicates that in addition to ATP, synthesis of other metabolites, specifically derived from glucose and glutamine, are critical for MuSC activation. Indeed, isotopic labeling in MuSCs using ^**13**^C_**6**_-D-glucose has shown that a significant level of histone acetylation marks derive from glucose-derived acetyl-coA ^2^. The mitochondrial pyruvate dehydrogenase (PDH) converts glucose-derived pyruvate to acetyl-coA. Pharmacologic suppression of PDH kinase, the negative regulator of PDH, correlates with increases in glucose-derived histone acetylation in MuSCs. This suggests that glucose is an important source of epigenetic modification during MuSC activation. Glutamine also supports cellular functions outside of oxidative phosphorylation. Among the non-respiratory roles of glutamine include serving as a precursor for synthesis of non-essential amino acids (NEAAs), as well as nucleotides ^22^. Radiolabeling experiments in C2C12 myoblasts show that glutamine predominantly contributes to protein biomass through generation of NEAAs ^23^. Glutamine metabolism has also been analyzed in other tissue resident stem cells, such as hematopoietic progenitor cells (HPCs). Analysis of glutamine-derived metabolites using high performance liquid chromatography mass spectrometry shows significant contribution of glutamine to nucleotide biosynthesis in CD34+ HPCs, and that this contribution is abolished following suppression of glutaminolysis ^24^. Inhibition of glutaminolysis as well as downregulation of the ASCT2 glutamine transporter in HPCs contributes to defects in erythroid differentiation and bias towards myeloid lineage commitment, suggesting that glutamine metabolism is necessary for balanced hematopoiesis. Addition of nucleosides to glutaminolysis-deficient HPCs restores erythropoiesis; it is possible that nucleoside supplementation to BPTES-treated MuSCs will rescue activation. Given these findings, MuSCs may metabolize glucose and glutamine not only to maintain bioenergetic homeostasis, but also to support epigenetic regulation of MuSC activation and biosynthesis of macromolecules. Further work is needed to dissect the diverse fates of glucose and glutamine in MuSCs during activation.

## METHODS

### Mice

All experiments were performed on samples derived from 3.5 to 7 months old male C57Bl6j mice obtained from Charles River. Mice were housed in the fully AAALAC-accredited animal care facility in the basement of the Eli and Edythe Broad Center for Regenerative Medicine and Stem Cell Research at USC. Animal use protocols were reviewed and approved by the USC Institutional Animal Care and Use Committee (IACUC).

### MuSC isolation and purification

MuSCs were isolated as previously described ^25^. Immediately after CO_**2**_ euthanasia, hind limb skeletal muscles were extracted and minced. Minced skeletal muscles were enzymatically digested into a single-cell solution by collagenase and dispase. The digested skeletal muscle sample was stained with CD31-FITC (BioLegend) CD45-FITC (BioLegend), Sca-1-PerCP (BioLegend), VCAM-PeCy7 (BD Biosciences), and Integrin α7-PE (Thermo Fischer Scientific). MuSCs were purified on the BD FACSAria IIu as a population of CD31-, CD45-, Sca1-, VCAM+, and Integrin-α7+ cells, and sorted into Ham’s F-10 plating medium (Cellgro) supplemented with 5 ng/mL basic fibroblast growth factor (Invitrogen), 10% FBS (Invitrogen), and 1X penicillin/streptomycin (GIBCO).

### Cell culture

Purified MuSCs in plating medium were seeded in 8-well chamber slides coated with ECM (Sigma E1270) over poly-D-lysine (Millipore) at 10,000 cells per well and allowed to adhere for approximately 1 hour post-isolation. After allowing MuSCs to adhere, plating medium was switched to growth medium, or Ham’s F10 media supplemented with 10% FBS and 10% horse serum. MuSCs were grown at 37 °C and 5% CO_**2**_ and growth medium was replenished every 4 to 12 hours. Freshly isolated MuSCs were imaged at 1 hpi for cell diameter analysis. MuSCs cultured for 24 and 40 hours were trypsinized then imaged for cell diameter measurement. To quantify protein synthesis, cells were cultured in normal growth medium then switched to cysteine- and methionine-free medium DMEM supplemented with 2 mM L-glutamine (VWR) and 100 µM HPG (Invitrogen), and cultured for 2 hours before fixation with 4% PFA. To measure entry into S-phase of the cell cycle, MuSCs were cultured in 5 µM EdU (Invitrogen) for 40 hours. HPG and EdU incorporation were detected using Alexa Fluor 488 azide. MuSCs were cultured in growth medium treated with 200 nM oligomycin (Cayman Chemical), 1 mM 2-deoxy-D-glucose (Sigma-Aldrich), 15 µM BPTES (Tocris), or 50 µM etomoxir (EMD Millipore) to block mitochondrial ATP synthesis, glycolysis, glutaminolysis, and fatty acid oxidation, respectively. The LIVE/DEAD Fixable Dead Cell Stain Kit (Invitrogen) was used according to the manufacturer’s protocol to detect cell viability under oligomycin treatment.

### Immunostaining

Immunocytochemistry staining was performed on MuSCs after fixation and permeabilization. MuSCs were fixed with 4% PFA for 10 minutes at room temperature then permeabilized using 0.3% Triton-X PBS for 15 minutes. Cells were then blocked with 10% donkey serum in 0.3% Triton-X PBS for 15 minutes. Cleaved caspase-3 was probed in MuSCs by incubating the cells overnight in 1:100 Cleaved Caspase-3 (Asp175) (5A1E) rabbit mAb (Cell Signaling Technology). MuSCs were stained for 3 hours with 1:500 donkey anti-rabbit Alexa 594 secondary antibody (Invitrogen) to detect cleaved caspase-3. pRB was stained for in MuSCs following EdU staining. MuSCs were first permeabilized then blocked in donkey serum for 20 minutes. After blocking, the cells were stained using 1:100 phospho-Rb (Ser807/811) (D20B12) XP rabbit mAb (Cell Signaling Technology). Secondary staining was performed using 1:500 donkey anti-rabbit Alexa 488 antibody (Invitrogen). All primary and secondary antibodies were diluted in 10% donkey serum.

### Time-lapse microscopy

MuSCs were plated on 8-well chamber slides coated with ECM and cultured for 90 hours at 37 °C and 5% CO_**2**_. MuSCs were then transferred to a temperature- and CO_**2**_-controlled chamber mounted on the microscope stage of the Zeiss Axio Observer Z1. Bright-field images were acquired every 10 minutes and cells were visualized using ZEN Black software. Time taken to complete the first division following initiation of image acquisition was only analyzed for MuSCs that remained in acquisition field.

### Extracellular flux assay

One day prior to the assay, Seahorse XF sensor cartridges were hydrated in XF Calibrant at 37 °C and 0% CO_**2**_ overnight. On the day of the assay, MuSCs were purified into plating medium and seeded immediately following isolation in 8-well Seahorse mini-plates coated with ECM. Plating medium was changed to growth medium treated with or without 15 µM BPTES and 50 µM etomoxir for MuSCs cultured for 24 and 48 hours at 37 °C and 5% CO_**2**_. For freshly isolated (FI) MuSCs, plating medium was changed to Seahorse assay medium (Buffered XF DMEM base medium containing 5 mM HEPES, 2 mM L-glutamine, and 10 mM glucose) after MuSCs were seeded. MuSCs were incubated in Seahorse assay medium for 1 hour at 37 °C and 0% CO_**2**_, then stained with propidium iodide and imaged to detect viable cells immediately before loading in the XF analyzer. MuSCs were administered 1 µM oligomycin and 0.25 µM rotenone and antimycin A during the assay. For measuring glycolytic metabolism, MuSCs were injected with 50 mM 2-DG before oligomycin and rotenone and antimycin A. ATP production rates were calculated from OCR based on the calculations described in Quantifying Cellular ATP Production Rate Using Agilent Seahorse XF Technology. Glucose-linked mitochondrial ATP production rate was calculated as the difference between baseline ATP production rate and ATP production post-2-DG injection. Glutamine or fatty acid-linked ATP production rate was the difference between baseline ATP production rate in control and BPTES- or etomoxir-treated MuSCs.

### Data analysis

All data are presented as mean ± standard deviation. For experiments that involved treatment of MuSCs with pharmacological inhibitors, MuSC population isolated from one mouse was split into control (untreated) and experimental (treated) groups post-isolation so that measurements can be taken in parallel. Time to the first division data are presented as the cumulative proportion of cells that have divided starting at 30 hpi and at every 10-hour interval afterwards. Cell migration was analyzed starting at 18 hpi and the distance traveled was measured at 2-hour intervals for the next 20 hours. Migration speed was calculated by dividing the distance traveled during each 2-hour interval by 2 hours; the speed is plotted as the speed at the midpoint of each time interval. Changes in MuSC area were analyzed starting at 18 hpi and measurements were taken every 6 hours, up to 42 hpi.

Excluding Figures 1E-F and 3A-C, all statistical comparisons were performed using a two-tailed, unpaired Student’s t-test; variance was tested using the F-test. A two-tailed, paired Student’s t-test was used for Figures 1E-F and 3A-C.

## ACKNOWLEDGEMENTS

This work was supported by grants from the NIH (R00AG041764), AFAR (Junior Faculty Grant), and The Baxter Family Foundation (to JTR).

## AUTHOR CONTRIBUTIONS

S.A. and J.T.R. designed the experiments. S.A., M.H.R., M.E., R.T., A.C., and J.T.R. performed and analyzed data for all experiments. S.A. and J.T.R. wrote the paper.

## FIGURE LEGENDS

### Supplemental Video 1

Movie generated from time lapse microscopy imaging showing the increases in MuSC cell size and motility in the first 48 hours after isolation. In the first 24 hours, the cells are small and display little migration. After 24 hours, MuSCs display a tremendous increase in cell size and movement. The video begins at 1.5 hours after FACs isolation and plating, each frame is 10 minutes apart, the time from isolation is displayed in the upper left (days:hours:minutes:seconds.ms).

